# A Spatial Proteomic Atlas of Tertiary Lymphoid Structures in Non-Small Cell Lung Cancer Identifies a Novel Predictive Class of Lymphoid Aggregates

**DOI:** 10.64898/2026.05.14.723890

**Authors:** Tyler Risom, Rajiv Jesudason, Evan Liu, Andrew Hill, Niha Beig, Conrad Foo, Ouida Liu, Eloisa Fuentes, Lisa Tai, Kedar Prasad, Jennifer Giltnane, Robert J. Johnston, Lisa M. McGinnis

## Abstract

Tertiary lymphoid structures (TLS) predict benefit from immune checkpoint inhibitors (CPIs), yet mature, germinal-center-rich TLS are infrequent in solid tumors by histological review. Here, using 38-plex MIBI spatial proteomics across 165 lymphoid structures from 14 NSCLC resections, we establish a continuum of TLS maturity using high dimensional compositional, spatial, and molecular features. We demonstrate that histologically-defined lymphoid aggregates (LA) comprise a heterogeneous class of structures, which span this continuum of maturity. We identify a subset of lymphoid aggregates that harbor follicular dendritic cell networks, T follicular helper cells, and activated B cell states characteristic of mature TLS, yet are not readily distinguished from other LA structures in our histological review. We developed a novel digital pathology classifier to identify mature LAs in CPI trials, and demonstrate in a retrospective analysis of Atezolizumab in advanced NSCLC that the inclusion of mature LAs greatly expands the biomarker-eligible population while maintaining strong predicted benefit. Together, these data redefine the biological spectrum of tumor-associated lymphoid aggregates and provide a framework for implementing maturity-informed TLS biomarker strategies.

## Introduction

Tertiary lymphoid structures are ectopic B cell follicles that form at the tissue site of chronic inflammatory diseases including cancer, and represent the culmination of the recruitment, spatial organization, and functional specialization of B and T cells^1–6^. TLS are defined by their similarity to B cell follicles in secondary lymphoid organs, where a germinal center (GC) reaction resides in the center of the follicle, hosting clonogenic B cell formation and supporting a proinflammatory niche in the surrounding tissue. In line with this function, the detection of TLS in tumor tissues by transcriptomic signatures^7–10^ or histological detection^11–13^ has been associated with positive patient outcomes across a myriad of solid tumor indications^5,14–16^, and is predictive of CPI benefit compared to standard of care^8,9,11,13^, including in our own non-small cell lung cancer (NSCLC) trials of the PDL1 inhibitor Atezolizumab^12^. However, the observation of archetypical TLSs with a large germinal center (GC) clearings appears to be indication specific, while commonly observed in melanoma^11^ these “mature” TLS representing the apex of a developmental trajectory are vastly outnumbered by immature lymphoid structures in NSCLC^7,12,17^, sarcoma^9^, hepatocellular^14,18^, renal^19^ and colorectal^16^ carcinoma, and the question remains of whether these immature states similarly impact a patient’s prognosis.

The impact of TLSs as predictive biomarkers is challenged by a lack of uniformity in the criteria defining the observed lymphoid structure states in tumors. While alignment around the definition of a germinal-center-positive mature TLS (TLSm, GC^+^ TLS, or secondary follicle-like TLS) is observed in the field, notable differences exist in the definition the less mature states of tumor lymphoid structures, including immature TLS (TLSi, primary follicle-like TLS) commonly defined by a small, amorphous germinal center clearing, and lymphoid aggregates (LA, GC^-^ TLS, early TLS) which lack an observable germinal center clearing but are identifiable as dense spatial aggregate of lymphocytes^15,20,21^. These criteria focus on the histological observation of germinal center clearing, rather than on the presence of anti-tumor effector cells central to proposed mechanisms of TLS-mediated anti-tumor immunity^2,3,22–26^. While key TLS effector cell presence is inarguably enriched in the germinal center^24,26,30^, the requirement of GC visual detection ignores amorphous or atypical arrangement of these cells in lymphoid structures, including cases where the GC edge is a few µm deeper into the block. Indeed evidence of false-negative TLSm detection is well documented in histological studies^27,28^ where the plane of tissue sectioning avoids sampling the GC and reveals a GC-negative lymphoid aggregate by histology, whereupon further sectioning into the block the GC is revealed. In this simple example, a ground truth TLSm exists across all levels, but histological sampling artifact drove structure misclassification. As germinal center reactions in TLS and SLO follicles are dynamic processes, initiating after antigenic challenge, persisting for variable lifespans, then resolving^24,29^, current TLS histological classification criteria attempt to bin a continuous process into distinct groups and can not readily separate true primordial GC-negative lymphoid aggregates from artifactual GC-negative structures. In these ways, the current over-reliance on GC observation in TLS biomarker strategy limits potential patient eligibility for transformative care where a GC^+^ TLS is not observed in a single 2-dimensional sampling of their disease.

To address these knowledge gaps and technical challenges and to improve upon how we leverage TLS and their immature structural states as biomarkers, here we performed paired 38-plex MIBIscope spatial proteomics profiling with histological lymphoid structure classification by a team of pathologists in a cohort of lymphoid-structure-positive, CPI-naive, NSCLC tumor resections samples. This cohort yielded 165 tumor lymphoid structures spanning the TLS maturity spectrum with >700 image based features extracted per structure, comprising a new NSCLC TLS atlas resource. By comparing histological definitions to unsupervised data-driven methods, we expand our understanding of the compositional and molecular heterogeneity within lymphoid structure classes, and reveal a novel subset of lymphoid aggregates that contain phenotypically mature B cells, follicular dendritic cell (FDC) processes, and other mature lymphoid structure features previously considered specific to GC+ TLS. Despite these characteristics identified in the MIBI profiles, these “mature LAs” were similarly classified by pathologist H&E review as the other LAs that lacked these mature features. This finding highlights the potential for the TLS field to better leverage immature lymphoid structure subsets for biomarker strategy improvements. Indeed we demonstrate in a retrospective trial of our PD-L1 inhibitor Atezolizumab that the quantification of these structures, in addition to TLS, doubled the size of the biomarker eligible patient population predictive of CPI benefit, as compared to histological detection of mature TLS. These findings highlight the meaningful compositional, spatial, and molecular maturity that can underlie germinal center negative lymphoid structures, and provides a novel biomarker strategy that identifies patients with mature lymphoid structures agnostic of their histological class, demonstrating how such structures may be leveraged to predict CPI benefit across solid tumor indications.

## Results

To illuminate the biological features across the spectrum of lymphoid structures in NSCLC and better understand influential processes present in immature states, we first generated a compositional and spatial atlas of 165 lymphoid structures across a cohort of primary, CPI-naive, lymphoid structure-positive NSCLC tumor resections (Fig1a, Table S1, N=14). Each sample was sectioned onto a specialized MIBIscope slide, with serial sections above and below the MIBI slide stained with hematoxylin and eosin (H&E) and reviewed by a team of 5 pathologists for exhaustive annotation of lymphoid structures according to a shared annotation protocol^17^, identifying 84 lymphoid aggregates (LA) as germinal-center-negative, lymphocyte-rich spatial aggregates; 17 immature TLS (TLSi) as lymphocytic aggregates with the observation of a small, irregular germinal clearing, measuring 200–750μm in diameter; and 14 mature TLS (TLSm) with pronounced germinal center clearing (Fig.1b, Table S2). A stromal infiltrate (SI, N=50) class of diffuse to dense lymphocytic infiltrate was also allowed as an annotation class upon final review, with additional SI regions being supplemented by a sixth pathologist. Each lymphoid structure class was assigned a numerical score according to common assumptions of their hierarchy along the TLS maturity axis, with SI=1, LA=2, TLSi=3, TLSm=4, enabling the calculation of a mean maturity score (MMS) across all pathologists across both H&E levels. The standard deviation between pathologists was also determined for each structure across each slide (SD-top, SD-bottom), and a global standard deviation (SD-global) using the average of slide level SDs, as well as a SD-Z metric taking the difference between slide-level maturity scores to evaluate the change in maturity scores across Z space. The MMS for each structure was used to assign a histological class label for comparative analysis, using the consensus class score +/- 0.5: SI (MMS = <1.49), LA (MMS = 1.5-2.49), TLSi (MMS = 2.5-3.49) and TLSm (MMS = 3.5-4) (Fig.1c). LAs were most prevalent, and comprised 75% of the aggregate structures in the cohort. TLSi showed the poorest concordance across pathologists and the largest score changes across H&E levels (largest global SD and SD-Z), potentially attributable to the small, nonuniform germinal center in this class that is prone to individual interpretation and change in Z space (Fig.1d).

The MIBI slide was stained with a 38-plex panel of metal conjugated antibodies targeting key immune, stromal, and TLS specialized cell biology (Fig.1e, Table S3). A 800x800um FOV was collected around each pathologist-identified structure, capturing the structure and its surrounding tumor microenvironment (TME, Fig.1f). We deployed a diverse set of analyses on each lymphoid structure (Fig.1g) including cell compositional analysis, pairwise cell-cell spatial relationships, multicellular niche identification, and classification and masking of the aggregate, GC, FDC network, & HEV objects in the images. In total we identified 24 cell types (Fig.1h) and 38 total phenotypic subsets of these cells (Fig. S1).

We next evaluated the compositional, spatial, and structural features between the four histological classes assigned by pathologist H&E review. Representative examples of each class are shown in Figure 2a, highlighting key TLS lineage marker expression, H&E appearance (Fig. 2b), architectural object detection, and cell type identities (Fig. 2c). Compositionally, SI regions consisted predominantly of T cells with variable B cell density, and showed the highest epithelial cell density of the classes as they were often annotated at the epithelial-stromal interface of tumor nests, and accordingly showed the highest PDL1+ fraction in their neighboring epithelium, indicative of tumor cell adjacency (Fig. 2d, S2a, b). In contrast, TLSi and TLSm regions were often observed at the tumor border, and resultantly showed minimal total epithelial cell density, and a high fraction of LAMP3-positivity in adjacent epithelial cells, consistent with normal lung pneumocyte mixing at the the tumor edge. LA, TLSi, and TLSm were all dense lymphocytic aggregates containing an increasing amount of B cells with high spatial colocalization to other B cells and T cells, all of which peaked in TLSm regions (Fig. S2c). Specialized cell types of the germinal center reaction, including follicular dendritic cells (FDC), T follicular helper cells (Tfh) and tingible body macrophages (TBM), were enriched in TLS and highest in TLSm structures, being sparsely observed in LAs (Fig. S2d).

**Figure 1:**
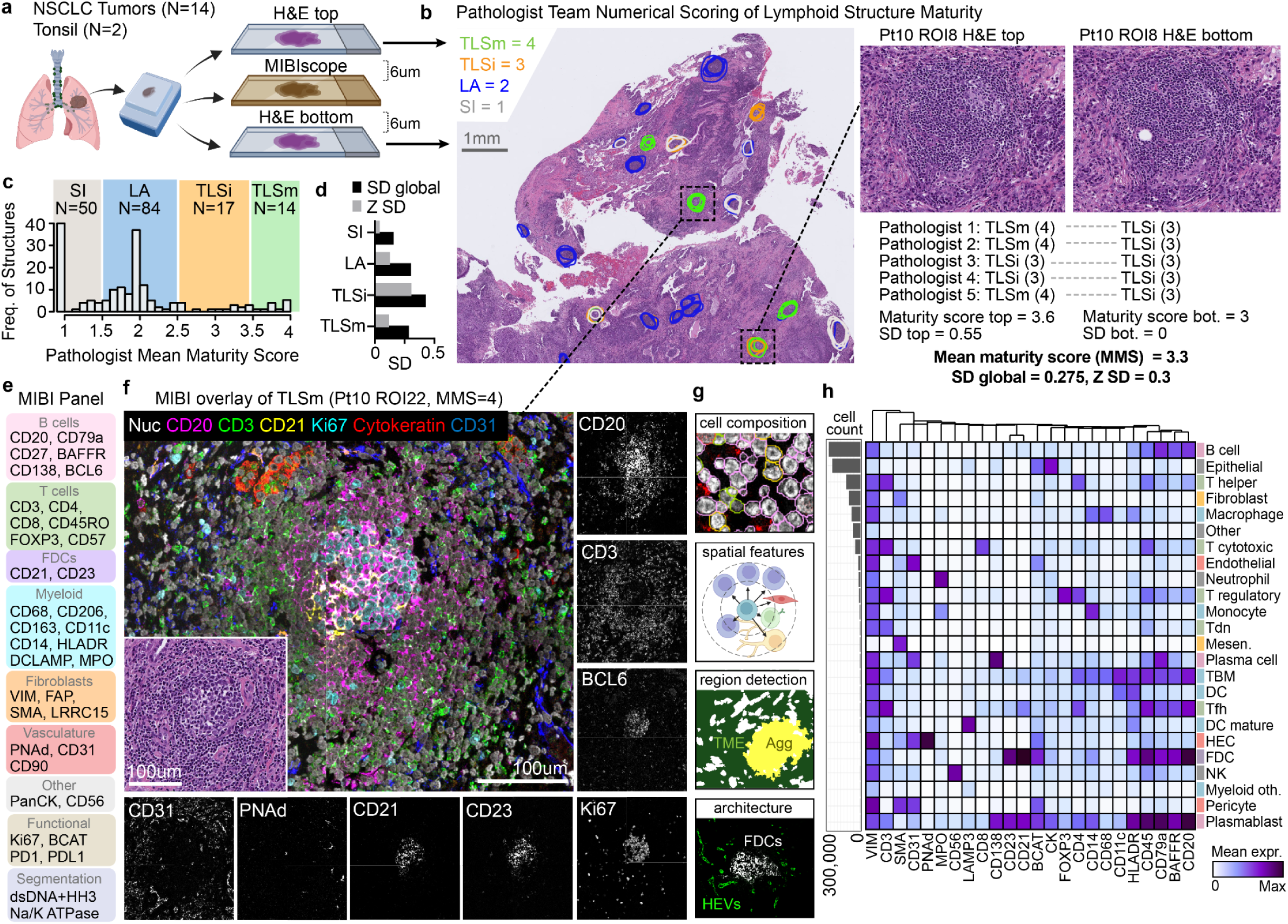
Overview of cohort and multiomic approach to lymphoid structure profiling. **a**. Schematic showing patient cohort and sectioning schedule. **b.** Overview of histological classification of SI (gray), LA (blue), TLSi (orange), and TLSm (green) regions by the pathologist team. An example H&E crop from both H&E levels is shown for a TLSi structure, with accompanying maturity score calculations using the pathologist annotations to identify the MMS and Z SD. **c.** Histogram showing the distribution of the 165 tumor lymphoid structures by mean maturity score, colors denote the bounds of each pathological class assignment for SI, LA, TLSi, and TLSm structures. **d.** Bar plot showing the global standard deviation across the two H&E slides (SD global), and the MMS SD between the top and bottom slide (SD-Z) for each class. **d.** Schematic showing the 38 markers in the MIBIscope NSCLC TLS panel. **e.** Schematic of the protein targets of the 38-plex MIBI panel used in the study, markers are organized into cell lineage themes. **f.** Representative MIBI overlay of a TLSm region showing nuclei (white), CD20 (pink), CD3 (green), CD21 (yellow), Ki67 (cyan), Cytokeratin (red), CD31 (blue), scale bar = 100um. Grayscale single marker abundance shown around the overlay, including additional markers BCL6, CD23, and PNAd. **g.** Schematic of analytical modules used in MIBI image analysis. **h.** Heatmap and bar plot showing the marker expression and abundance of the 24 cell types profiled in the study.

**Figure 2:**
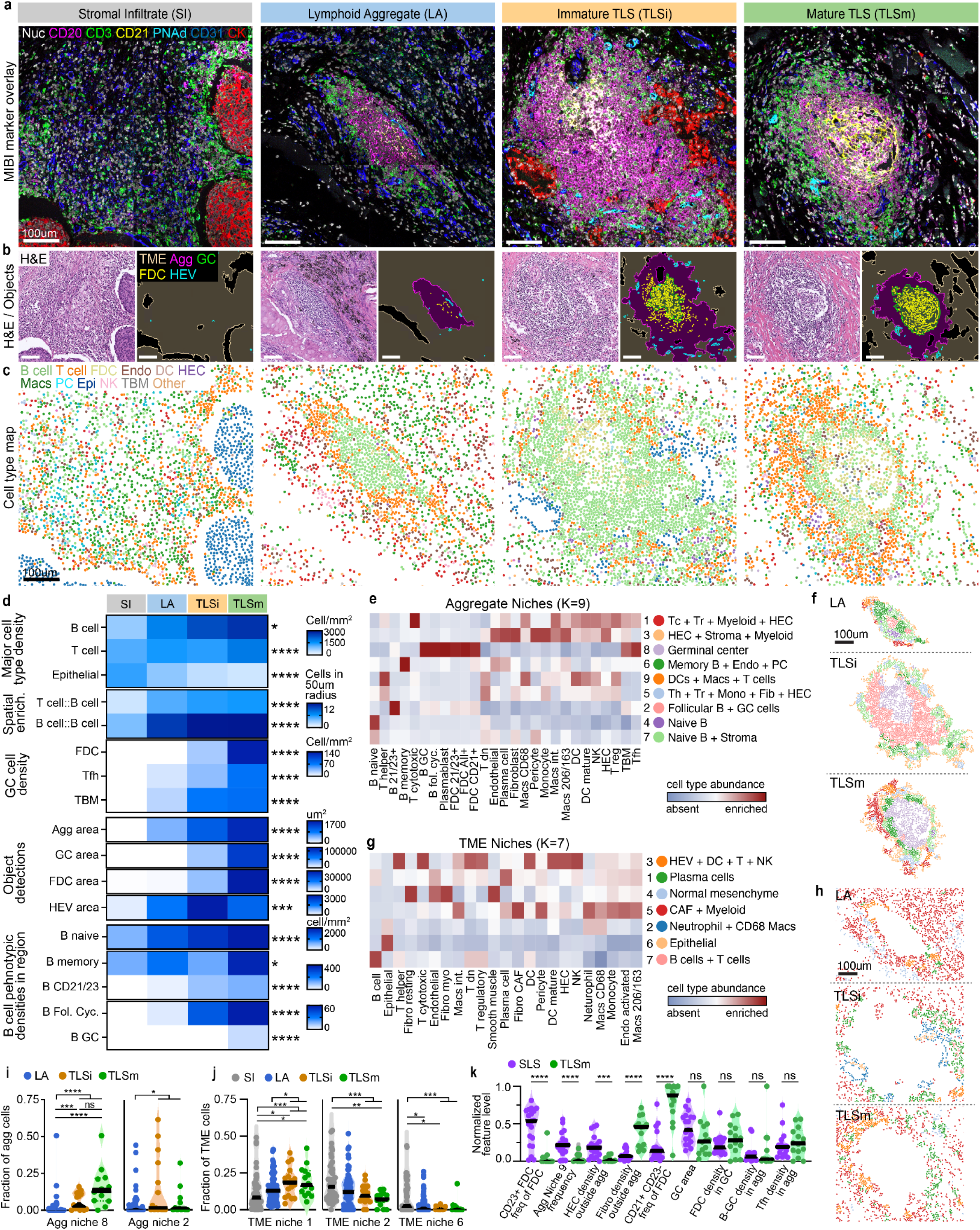
A NSCLC lymphoid structure atlas reveals class defining features. **a**. Representative MIBI marker overlays for each pathologist class showing CD20 (magenta), CD3 (green), CD21 (yellow), PNAd (cyan), CD31 (blue) and pan-cytokeratin (CK, red), scale bars = 100um. **b**. H&E images (left) and pixel classifier detected object maps (right) for the corresponding structure shown in a, scale bars = 100um. **c.** Cell type map colored by the 16 major cell type identities, scale bar = 100um**. d.** Heatmap showing the median values of cell type density, spatial enrichment of major cell types, specialized GC cell type density, areas of detected objects, and the density of B cell subsets across the whole 800x800 image ROI, between the four pathologist classes. Asterisks denote significant differences by Kuskal-Wallis test between all groups * P < 0.05, ** P < 0.01, *** P < 0.001, **** P < 0.0001. **e.** Heatmap showing the cell type enrichment in the 9 identified aggregate niches, with cluster ID, color identifier, and descriptive name of cluster shown to the right. **f.** Cell map of intra-aggregate cells in representative LA, TLSi, and TLSm, colored by niche identity from *e*, scale bar = 100um. **g.** Heatmap showing the cell type enrichment in the 7 identified TME niches, cluster ID, color identifier, and descriptive name of cluster shown to the right. **h.** Cell map of TME cells in representative LA, TLSi, and TLSm, colored by niche identity from *g*, scale bar = 100um. **i-j.** Violin plots showing the frequencies of select aggregate niches (i) or TME niches (j) across the four pathologist classes. Asterisks denote significance by Kruskal Wallis test with Dunn’s multiple hypothesis correction: * P < 0.05, ** P < 0.01, *** P < 0.001, **** P < 0.0001. **k.** Violin plot comparing features between TLSm (N=14) and tonsil SLS (N=19), asterisks denote significance by Mann-Whitney test: *** P < 0.001, **** P < 0.0001.

Correlated with these compositional signatures were changes in the architecture of the lymphoid structures, revealed by the output of deep learning pixel classifiers trained to detect lymphoid aggregates, germinal centers, the network of FDC processes, and high endothelial venules (HEVs). Larger aggregates were detected in TLSi vs LA, and TLSm vs TLSi, and germinal centers were predominantly detected in TLSi and all TLSm, with mature TLS having a median GC size twice that of TLSi (Fig. S2e). Similarly, the FDC network area in TLSm was double that of TLSi. To our surprise, however, HEV classifier detections showed a similar area across LA, TLSi and TLSm regions, potentially related to their proposed early establishment in the TLS development process, though it is important to acknowledge that the increased aggregate sizes in TLSi and TLSm restricts the sampled TME area for potential HEV detection. Consistent with their germinal center positivity, TLSi and TLSm were enriched in phenotypically mature B cell subsets including memory B cells, follicular-like B cells coexpressing CD21, CD23, CD27, HLA-DR, and PD1 (annotated as B_CD21/CD23), and their Ki67^+^ “follicular cycling” counterpart (Fig. S2f). Interestingly, TLSm were uniquely enriched in BCL6+ “B-GC” cells, and normalization of GC B cell density to GC area further distinguished TLSm from TLSi (Fig. S2g).

To better understand the spatial units that comprise TLS and their less mature counterparts, as well as their proximal tumor microenvironment (TME), we identified multicellular niches of spatially co-enriched cell types within two distinct histological compartments: first restricted to cells within the aggregate boundary, and second restricted to cells outside of the aggregate boundary in the surrounding TME (Fig. S2h). Within the aggregate compartment we identified 9 niches, including niche 8 which showed specificity to the center of TLSi and TLSm structures (Fig. 2e, f). Interrogation of cell type composition in niches revealed that aggregate niche 8 was enriched for B-GC cells, Tfh, FDCs and TBM, likely identifying the cellular interplay within the germinal center reaction, and accordingly, we found TLS structures to be enriched in niche 8 compared to LA. Often bordering this GC-specific niche was aggregate niche 2, composed of CD21/CD23+ follicular B cells and a sampling of other specialized GC cells. Niche 2 levels were highest in TLSi, and showed enrichment in TLSs compared to LAs (Fig. 2i). Outside of the aggregate we identified 7 TME niches, including those describing TME composition around tumor cells (TME niche 6) and neutrophils (TME niche 2) that were enriched in SI regions (Fig. 2j). A plasma cell rich TME niche 1 was also identified that showed enrichment in TLS vs. SI and LA regions.

Finally, we compared compositional, spatial, and architectural features between TLSm (N=14) and germinal-center-positive secondary lymphoid structures (SLS) from tonsil (N=19) to explore how tumor TLS adhere to evolved processes established in secondary lymphoid organs. Compared to TLSm, SLSs showed a higher frequency of CD23 positivity in FDCs, and an increased presence of aggregate niche 9, describing the spatial colocalization of dendritic cells and T cells in the aggregate (Fig. 2k). In addition, the cellular composition outside of the aggregate was distinctive, but reflected an expected pattern of tumor microenvironment cells surrounding TLSm, including enrichment of fibroblasts, monocytes and macrophages, whereas B cells, T cells, and HEC density was enriched around tonsil SLS. To our surprise, beyond these FDC phenotypic differences, TLSm and SLS showed similar germinal center features with no significant differences observed regarding germinal center size, FDC density in the GC, B-GC, B fol. Cyc., plasmablast, Tfh, and TBM density in the aggregate, and the frequency of aggregate niche 8 which describes the multicellular interactions in the GC (Fig. S2j, k).

Taken together, this compositional and spatial atlas of tumor lymphoid structures establishes definitive features of the major histological classes, identifying new discriminators between immature and mature TLS, and largely reinforcing that germinal-center-related features are central to the identification and classification of TLS structures from lymphoid aggregates.

Because LAs lack the hallmark germinal center used for histological classification, they have been treated as a heterogeneous, poorly-defined class in TLS studies, despite their prevalence in tumors. We were excited to leverage these data to explore the molecular and cellular heterogeneity in LAs, and applied the Uniform Manifold Approximation and Projection (UMAP) analysis to all cohort structures leveraging 759 structure-level features to address their interrelationships in the context of known maturity classes. We subsequently performed Slingshot analysis^31^ on the projection, a tool used for evaluating cell lineages and pseudotimes in single cell expression experiments, which yielded a continuous value for the “pseudotime” of each structure, termed here the continuum location (CL), in addition to generating 5 similarity clusters along the CL axis (Fig.3a-c, S3a-c). SLS regions were well separated from the tumor lymphoid structures with the highest CL values, potentially related to their unique tissue microenvironmental features, and the SLS exclusively comprised the most advanced CL cluster K5 (Fig. 3d). TLSm and TLSi structures were predominantly organized together with high CL values and clustered in the most advanced non-SLS clusters, K4 and K3. Conversely, the SI class regions showed the lowest CL values and comprised the majority of the least advanced CL cluster K1. The LA class, however, showed marked heterogeneity, spanning all non-SLS clusters (K1–4) and the entire CL range from SI to TLSm (Fig.3e,f). This drastic CL heterogeneity in LAs suggested their compositional, spatial, and molecular identity approximates SLS in the most mature subset and SI in the least mature. We tested this hypothesis by training a linear discriminant analysis (LDA) classifier to classify SLS from SI regions, and deployed the classifier on all cohort structures (Fig. S3d). The results showed that the LD1 variable was highly correlated to established maturity features (Fig. S3e) and reinforced the CL analysis: TLSm had the highest mean LD1 values, SI the lowest mean LD1, and LAs spanned the full range of LD1 (Fig. S3f), reinforcing the existence of profound compositional and spatial feature heterogeneity in this class.

**Figure 3:**
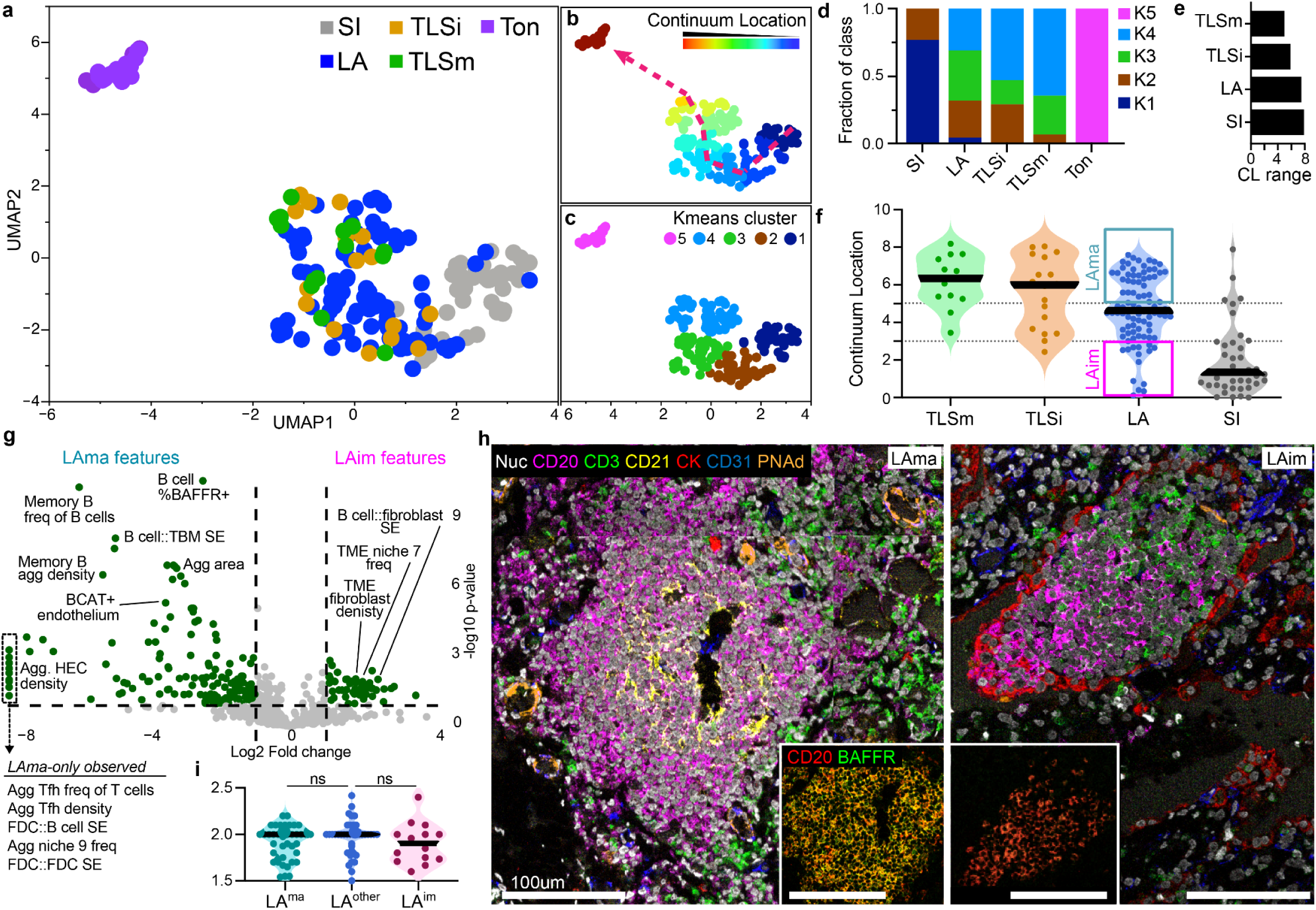
Unsupervised lymphoid structures analysis reveals a continuum with distinct maturity subsets of LAs. **a**. Scatterplot showing the UMAP arrangement of lymphoid structures based on their high-dimensional relationships between 580 feature inputs, color denotes pathologist class or tonsil SLS (violet). **b.** The CL value of each lymphoid structure is denoted by color overlay on the scatterplot. A red line is overlaid to show the direction of the CL values from low to high CL. **c.** Slingshot k-means cluster identity is overlaid on the points by color. **d.** Area column plot showing the fraction of slingshot k-means clusters within each pathologist class. **e.** Bar plot showing the variance in slingshot trajectory CL within each pathologist class. **f.** Violin plot showing the slingshot trajectory CL distribution in each pathologist class. Boxes are drawn on the LA violin showing the subset identified as LA^ma^ and LA^im^. **g.** Volcano plot showing enriched features in LA^ma^ vs LA^im^ classes of LA structures. A set of features specifically enriched in LA^ma^ (“LAma-only observed”) are shown to the left, where log2fc = -infinity. **h.** MIBI overlays of 7 markers showing a representative LA^ma^ and LA^im^ region, scaled bars = 100um. **i**. Violin plot showing the mean maturity scores of LA^ma^, LA^im^, and non-LA^ma^/LA^im^ LA regions “LA^other^”.

To better understand the features driving the CL heterogeneity amongst LA structures, we evaluated the differential feature enrichment between LA structures with high CL values aligned with the most advanced TLS CL cluster (cluster 4, CL>5) termed mature LAs (LA^ma^), and LAs with CL aligned with the least mature CL clusters (cluster 2, 1, CL<3) termed immature LA (LA^im^). LA^ma^ showed selective enrichment of multiple features associated with mature TLS that were absent in LA^im^ structures, including features denoting FDC, TBM, and Tfh presence within the aggregate, FDC and B cell spatial enrichment, and agg-niche 9, describing DC, T and B cell interaction (Fig. 3g). In addition, LA^ma^ structures were larger than LA^im^, contained higher densities of memory B cells, and displayed higher frequencies of BAFFR+ and HLA-DR+ B cells. Visual inspection supported these findings and revealed the presence of FDC cells and FDC dendritic processes innervating LA^ma^ structures, as well as more proximal HEVs, and higher %BAFFR^+^ B cells (Fig. 3h). Despite these marked differences in the cellular composition, morphology, and phenotypic states, pathologist mean maturity scores showed no statistical difference between LA^ma^ and LA^im^ structures (Fig. 3i), indicating they show no preference for assignment as TLSi and are indistinguishable from other LAs given our review protocol.

In three-dimensional imaging studies, tertiary lymphoid structures have been shown to exist within networks of numerous lymphoid structures interconnected by HEV networks and lymphocyte trafficking routes. We sought to explore spatial relationships on a structure level to understand if LA^ma^ are spatially enriched near TLS compared to other LAs, potentially indicating their preference for mature lymphoid structure networks. To accomplish this, we generated graph networks of LA, TLSi, and TLSm structures that fell within a 2mm radius of one another, equivalent to approximately 5 TLS diameters. This analysis identified 20 multi-structure networks across 11 of 14 samples, with a mean of 5 lymphoid structures per network (Fig. S4a). In larger samples, such as patients 9 & 10, numerous networks are observed including those that are restricted to LAs, and those containing TLS (Fig. 4a,b). We first investigated whether TLSm presence correlated with network size or connectivity. TLSm-containing networks (7 of 20) exhibited higher mean node and edge numbers (Fig. 4c,d). In addition, TLSm^+^ networks also contained the majority of TLSi structures (Fig. 4e). LA^ma^ structures showed a distribution similar to TLSi, with 75% located in TLSm-containing networks, and a smaller fraction of LA^ma^ existed outside of networks compared to LA^rest^ (Fig. S4b). Further, we explored the edge lengths connecting structures to query the pairwise spatial relationship between LA^ma^ and TLS, and found that LA^ma^ were significantly closer to their nearest TLS neighbor than LA^rest^ (Fig. 4f), and across the entire tissue section LA^ma^ showed smaller mean distances to any TLS compared to LA^rest^ (Fig. 4g). Together, these analyses suggested that enrichment of histologic maturity may exist on a network level, where increased connectivity and size correlate with enrichment of mature lymphoid structures, including TLS and LA^ma^.

**Figure 4:**
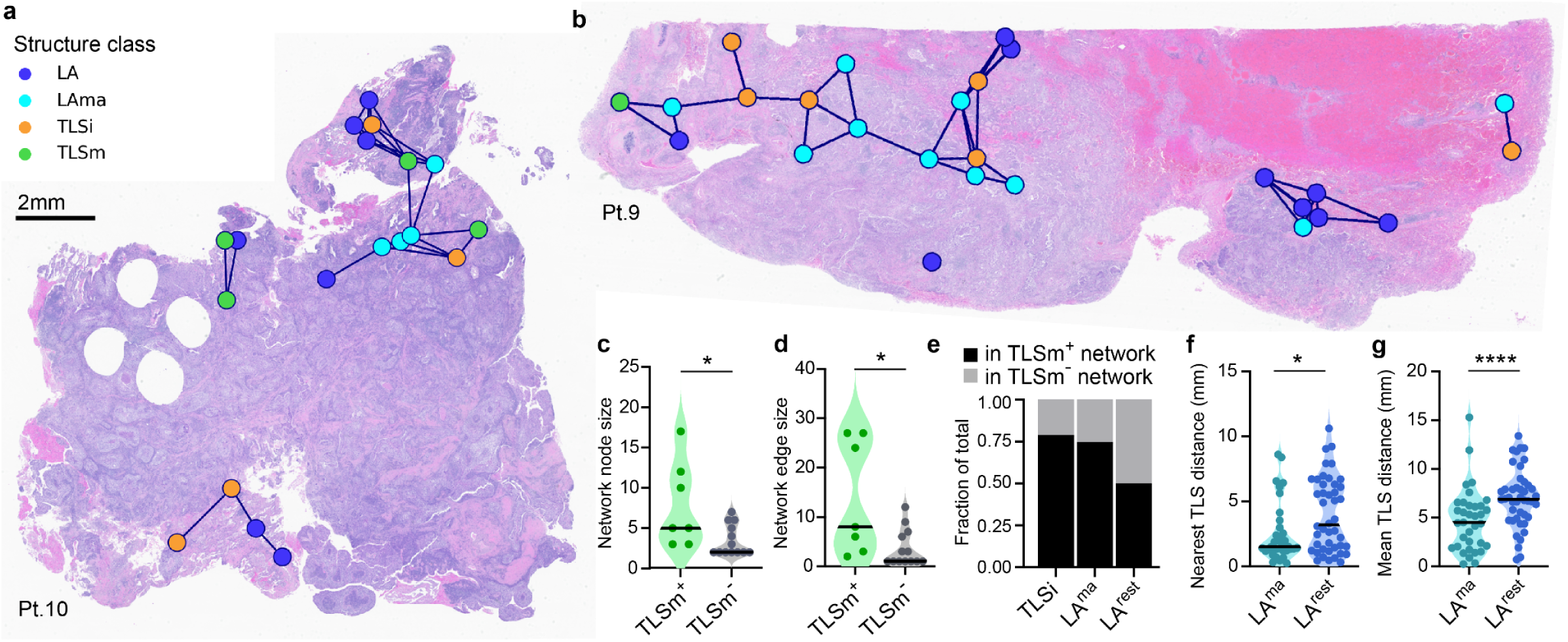
Mature lymphoid structures are enriched in large networks. **a-b**. H&E images of patient tumor resections overlaid with colored nodes identifying each LA (blue), LA^ma^ (cyan), TLSi (orange), and TLSm (green) structure in the sample. Structures within networks are connected by edges (black lines). **c-d.** Violin plot comparing the node size (c) and edge number (d) of TLSm-containing networks vs TLSm-negative networks. **e.** Area column plot showing the fraction of TLSi, LA^ma^, and LA^rest^ that exist in TLSm-containing vs. TLSm-negative networks. **f**. Violin plot showing the distance to the nearest TLS from LA^ma^ vs. LA^rest^. **g**. Violin plot showing the mean distance to all TLS from LA^ma^ vs. LA^rest^.

Given their compositional maturity and spatial enrichment with TLS, we found it paramount to understand if histological LA^ma^ detection is associated with positive patient outcomes. Importantly, LA^ma^ were observed at similar rates in TLS^+^ and TLS^-^ patients in this cohort (Fig. S5a), identifying a TLS-negative, LA^ma^-positive patient population that would be excluded from current TLS-centric biomarker strategies. To explore their relationship with outcomes, we first had to design a histological detection workflow. Understanding the lack of marked differences LA^ma^ and other LAs, we approached this through leveraging high resolution 40x scans of the H&Es with an in-house DINOv2-based vision transformer model that embedded 98x98px patches of a 882x882px crop surrounding each lymphoid structures (Fig. 5a, S5b). The resulting patch embeddings then underwent hierarchical K-means clustering (k = 128), and a random forest classifier was trained to differentiate “mature lymphoid structures” (MLS) including LA^ma^, TLSi or TLSm, from “immature structures” (LA^rest^ and SI) using the cluster composition of each region. We tested the accuracy of this classifier on two withheld patient samples that encompassed 29 MLS and 7 immature structures, and obtained a F1 score of 0.85 and precision of 0.91 in this in-cohort validation test (Fig. 5b, S5c). The decision tree had two decision points, both heavily weighing the absence of two cluster identities that showed specificity to non-MLS structures (Fig. S5d).

**Figure 5:**
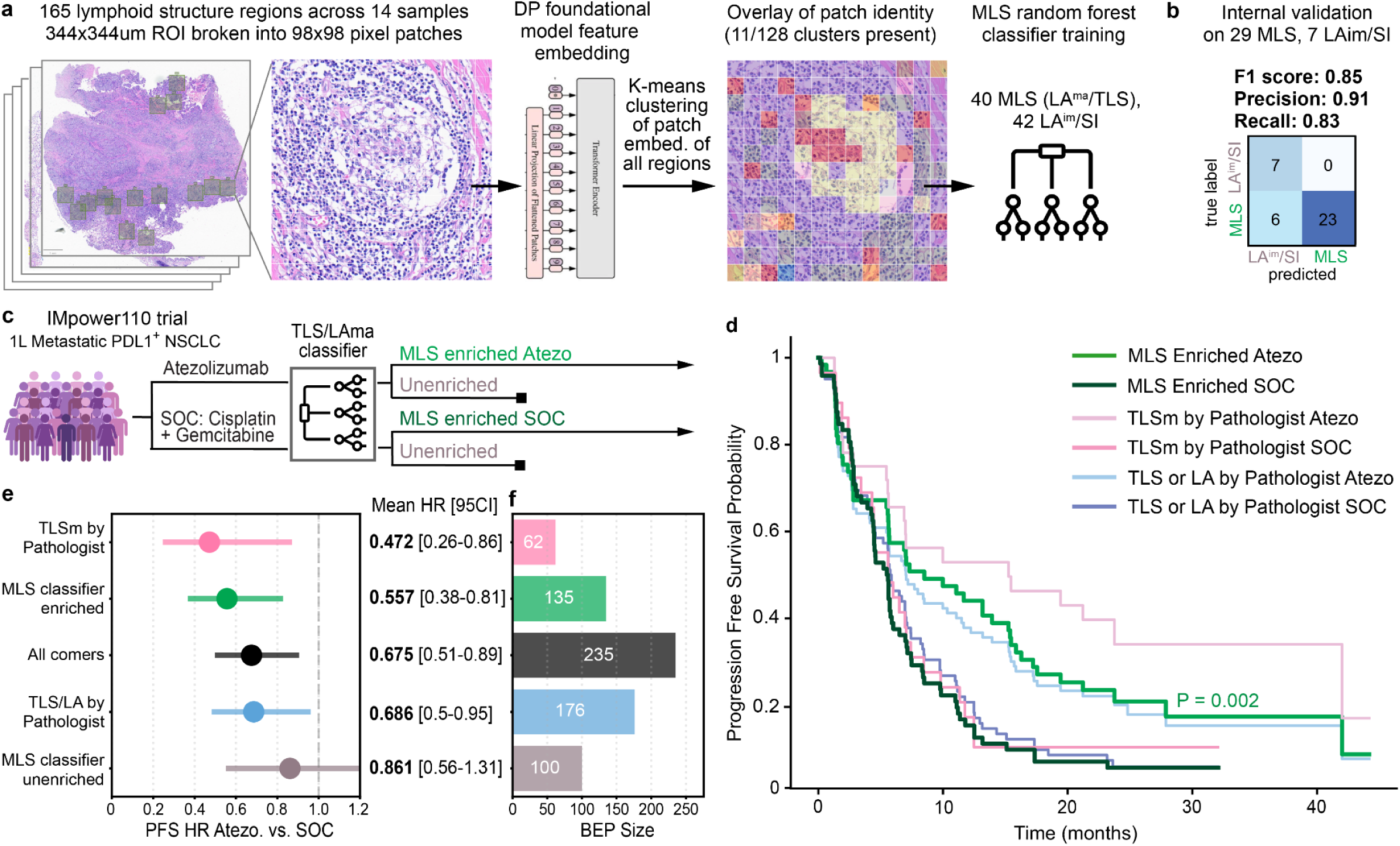
An novel biomarker strategy quantifying mature lymphoid structures predicts CPI benefit with increased BEP size. **a**. Schematic showing the workflow to train a LA^ma^/TLS patient sample classifier. A 882x882 pixel H&E crop centered on each of the 165 tumor lymphoid structures was broken into 98x98-pixel patches, and subsequently feature embeddings were generated for each patch with a foundational model. Patches were then clustered into 128 clusters using their feature embeddings, and a random forest classifier was trained to discriminate LA^ma^ & TLS “mature lymphoid structure” (MLS) regions from LA^im^ & SI regions by patch composition. **b.** The classifier was tested on 2 withheld images containing 29 MLS & 7 LA^im^ & SI samples, with performance metrics displayed. **c.** Schematic showing the different treatment arms of the IMpower110 trail and how the MLS classifier is used to separate MLS enriched and unenriched groups on both arms. **d.** Kaplan-meir (KM) plot showing the outcomes of patients in the MLS BEP on the Atezolizumab treatment arm (light green) versus standard of care chemotherapy (SOC, dark green), with P value comparing the two arms displayed. Alternative biomarker strategy KM plots are overlaid including TLSm detection by histology (pink & red), and TLS + LA detection by histology (light blue and blue). **e**. Forest plot showing the mean hazard ratio of patients in the different BEPs with 95% CI displayed with a line, mean HR and 95%CI values are also written to the right. **f**. Bar plot showing the number of patients in each BEP with N overlaid.

We then evaluated the classifier in an external H&E dataset to elucidate the associations of MLS to patient outcomes. We analyzed the IMpower110 trial where 235 patients with metastatic PDL1^+^ NSCLC were randomized into treatment arms of Atezolizumab or standard of care chemotherapy (Fig. 5c). In order to first locate lymphoid structures across the patient H&E images, we deployed a Detectron2 TLS instance-segmentation model trained to identify TLS-like structures, termed TLSv4.1^17^. TLSv4.1-detected regions that met a probability threshold of 0.5, identifying LA structures and more mature states (e.g. TLS), were cropped into 882x882px images and subsequently broken into 98x98px patches and embedded and classified using our MLS ViT-classifier workflow above. Patients were identified as “MLS-enriched” if one or more identified lymphoid structures was classified as a MLS. Kaplan meier analysis comparing patient progression free survival (PFS) in the biomarker-enriched population (BEP) showed strong PFS benefit in the Atezolizumab CPI-treatment arm versus standard of care chemotherapy (p=0.002, Fig. 5d).

To evaluate the impact of including LA^ma^ in the biomarker strategy, we compared MLS biomarker results against two classical TLS biomarker strategies based on manual pathologist review of the H&Es for either TLSm, or “all TLS & LA”. As expected and recently reported^12^, manual TLSm detection by histological review predicted CPI benefit in this trial, and this strategy slightly outperformed the MLS-classifier in its mean PFS hazard ratio (HR) of 0.472 vs. 0.557, respectively (Fig. 5e). However, comparison of BEP size showed that TLSm were histologically detected in only 62/235 patients, whereas the MLS-classifier identified nearly two thirds of the cohort as being biomarker enriched (135/235), signifying a doubling of the BEP size (Fig. 5f). The BEP difference included 13 patients identified as MLS^+^ that lacked detection of any TLS or LA by manual review, and 60 patients that lacked TLSm but contained other mature states like TLSi or LA^ma^ (Table S4). This resolution of the MLS classifier to separate LA^ma^ from the rest of LAs was meaningful, as the MLS classifier excluded 50 patients that were present in the “all TLS & LA” BEP, whose inclusion negatively impacted the predicted benefit, increasing the mean HR from 0.557 (MLS) to 0.686 (all TLS & LA). Poor outcomes in the MLS-unenriched BEP (mean HR = 0.861) further support this conclusion, which is worse than all-comers (mean HR = 0.675), indicating that immature forms of lymphoid aggregates do not support CPI responses in this setting. Taken together, these data suggest the presented MLS biomarker strategy represents an automated method to greatly expand the patient population predicted to benefit from CPI treatment, aided by machine learning detection of lymphoid structures that may be missed by manual review, but more so from separating the beneficial mature lymphoid structures beyond TLSm from their immature counterparts.

## Discussion

TLS are increasingly recognized as critical mediators of local immune responses within the tumor microenvironment, serving as sites of antigen presentation, lymphocyte activation, and coordination of adaptive immunity. As a whole, our results describe how the histological classification of TLS intersects with single cell, molecular, and spatial features of NSCLC lymphoid structures. This dataset serves as a valuable resource for clarifying lymphoid structure identity as the use of TLS in retrospective analysis, or as biomarkers in prospective patient selection, rapidly expands in the immuno-oncology field. In support of their rapidly evolving incorporation into therapeutic guidance, this work also contributes a novel machine learning workflow for the automated detection of mature lymphoid structure states in the H&E domain.

These data reiterate defining features of LAs, TLSi, and TLSm in the field, demonstrating an incremental increase in aggregate size, specialized germinal center cell type presence, and advanced B cell phenotypes amongst these lymphoid structure classes, and reinforces that detection of the germinal center is the salient differentiator amongst the classes. The high-resolution, hyperplex imaging data in this study offered a novel opportunity to parse germinal center detection from compositional biology, enabling a deeper dive into what germinal center features truly differentiate mature TLS from other states, including secondary lymphoid structures in the tonsil. Through leveraging digital masks of the GC compartment via deep learning pixel classifiers, we demonstrate that most GC specialized cell types are similar in prevalence across TLSi, TLSm and tonsil SLS when normalized to GC area, e.g. GC cell density, including FDCs, Tfh cells, TBM, and plasmablasts. Only the GC density of BCL6+ GC B cells significantly differed between TLSi and TLSm, and were similarly enriched in the SLS GC, neighbored by CD23-enriched FDCs indicative of robust light zone reactions.

The similarities in germinal center composition across TLS subsets suggests the overweighted use of germinal center area for structure classification may lead to misclassification of lymphoid structures that have similar effector cell presence and functional state. This risk is particularly relevant in the 2-dimensional study of these spherical/ovoid objects, where the sectioning on the edge of a mature TLS may appear as a 2-dimensional TLSi if minimal GC is sampled, or a LA if no GC is sampled in the sectioning plane. Indeed we observe such anecdotal cases of two-dimensional sampling artifact in this dataset, where a structure annotated as a lymphoid aggregate on one level appeared as a GC-rich TLSm approximately 7 μm deeper. Extensive serial sectioning or three-dimensional imaging of TLS-rich tumor specimens is needed to help resolve the prevalence of such false-positivity and -negativity in our histological TLS classification protocols.

Our unsupervised analysis of hundreds of structure-level features reconciles this GC overreliance in classification, revealing a continuum of lymphoid structure maturity rather than discrete categories. Our analysis of the resultant data provided a useful single statistic of continuum location for interrogating structure maturity. While median CL values supported the established hierarchy of histological classes of TLSm>TLSi>LA>SI^1,12,20–22,32^, the CL variance within classes was more striking and illuminated the independence of GC size in this unsupervised arrangement. Some GC^+^ TLSm exhibited low CL values, whereas many GC-negative LAs showed high CL values, guiding our subsequent identification of the CL-high LA^ma^ state. The LA^ma^ subset demonstrates the existence of mature cellular composition and molecular signatures independent of an observable GC clearing, including FDC cells and their process network, Tfh cells, TBM, and BAFFR^high^ B cells. Despite these marked compositional differences, these LA subsets appear indistinguishable given our H&E annotation criteria, with no LA^ma^ consensus-classified as TLSi on either H&E level, and all LA subsets having statistically insignificant differences in MMS pathologist maturity score. Our vision transformer workflow successfully classified LA^ma^ from LA^rest^, suggesting that morphological distinctions are learnable, but may benefit from higher magnification and refined H&E interpretation criteria.

Our biomarker strategy presented here prioritizes the detection of mature lymphoid structures, regardless of GC observation, and demonstrates the clinical impact of this approach. The addition of LA^ma^ to TLSm and TLSi detection greatly expanded the biomarker positive population while maintaining robust outcome predictions for Atezolizumab. 31% of patients lacked TLSm but harbored other molecularly mature lymphoid structures. For these patients, this biomarker expansion may enable access to effective immunotherapy that would have been previously withheld due to absence of GC^+^ TLS. In addition to this key advance, this biomarker strategy aligns with other efforts^17,33–36^ in a push towards automated digital pathology workflows for leveraging TLS as biomarkers, eliminating the access barriers and time considerations of pathologist histological review. Validating this approach outside of our Atezolizumab trials and in solid tumor indications beyond NSCLC will be key to optimizing this workflow strategy, and we are actively exploring these avenues. As clinical trials evaluating the prospective use of TLS for CPI patient selection are underway^37,38^ we are excited to refine this strategy for potential use in CPI trials in the near future.

## Methods

### Patient Cohort

The patient cohort consisted of 14 NSCLC primary tumor resections identified as high quality and having lymphoid aggregate presence by pathologist histological review, details on each sample are described in Table S1. FFPE tissue blocks were procured from BioreclamationIVT (BioIVT, Westbury, NY), ProteoGenex (Culver City, CA), Cureline Inc. (South San Francisco, CA), MT Group (Los Angeles, CA), or Discovery Life Sciences (Alameda, CA). Disease TNM staging ranged from T1N0M0(1A) to T4N1M0(3). Patient ages in the 11/14 with known status ranged from 57 to 72, 8/11 were male, and all patients with known race status were white. Patients were immunotherapy naive with mixed history of standard of care chemotherapy and radiation. 9/14 samples were diagnosed as squamous cell carcinoma.

### Tissue preparation

Tissue sectioning, histology, and MIBIscope staining were performed at Genentech in the histopathology core of the Research Pathology Department. Tissue blocks were sectioned at 5um thickness onto either superfrost plus slides for hematoxylin & eosin staining (MilliporeSigma, Burlington, MA) or onto to gold-and-tantalum sputtered MIBIslides (Ionpath Inc., Menlo Park, CA) and prepared according to vendor protocols. Briefly, MIBIslides were baked for 1hr at 70° C, deparaffinized in xylene, rehydrated in an alcohol series, and underwent heat induced epitope retrieval in Ph9 target retrieval buffer (Dako inc) in line with Angelo lab best practice protocols^39^. Slides were then blocked, and incubated overnight at 4deg C in a metal-conjugated antibody cocktail of 37 markers (Table S2). The following day the slides were washed and incubated with a second antibody cocktail containing anti-DsDNA/HH3-89Y and anti-biotin-160Dy for one hour at 4deg C. Slides were then washed, fixed in 2% glutaraldehyde (Electron Microscopy Sciences, Hatfield, PA), and dehydrated in an alcohol series. Slides were stored in a nitrogen cabinet prior to imaging.

### Pathologist histological annotation of lymphoid structures

A team of five board certified pathologists reviewed digital scans of the 28 NSCLC H&E slide images with the task of annotating lymphoid aggregates, immature TLS, and mature TLS according to a shared protocol. Whole slide screening was performed at 1-2x magnification to identify lymphoid collections, followed by confirmation and categorization at 4-8x magnification using standardized definitions of the structures. Lymph nodes and remote bronchial-associated lymphoid tissue (BALT) were excluded from screening as secondary lymphoid structures; however, BALT adjacent to the tumor was included. Annotations were not performed in areas with >20% tissue artifact. Standardized definitions included the following:

Lymphoid Aggregate (LA): LAs were identified as dense collections of small monotonous lymphocytes measuring 200-750µm with distinct borders at 1-2x magnification, not embedded in diffuse lymphoid background. At 4-8x magnification, LAs lacked germinal centers, though some high endothelial venules (HEV) may be present but typically not in high density.

Immature TLS (TLSi): TLSi were distinguished from LA by the presence of a mixture of small and large cells representing an early germinal center stage, though lacking complete germinal center definition. Structures measuring 200-750µm with central pallor and/or high density of HEV at 8x magnification were classified as TLSi. Alternatively, TLSi could be identified by high density of HEV without definite germinal center cells, provided a dense lymphoid aggregate was associated with the HEV and the structure was clearly organized rather than representing non-specific tissue background. TLSi demonstrated greater organization than simple lymphoid aggregates

Mature TLS (TLSm): TLSm were classified as lymphoid infiltrates of any size composed of tightly packed lymphocytes with a well-defined germinal center recognizable as central pallor at 1-8x magnification. The germinal center was characterized by distinct, well-defined borders with typical ovoid or round shape, though HEV may or may not be present. Each TLSm annotation contained only one germinal center; structures with two adjacent germinal centers were annotated separately.

To remove recency bias in review, the two H&E images per patient were maximally separated in review order. Once each pathologist had completed their annotations, annotations were merged into one image per slide for the team to review with the option to correct their original annotation, including to a (diffuse) stromal infiltrate (SI) class.

A sixth pathologist, blinded to the team review results, was used to place MIBIscope imaging locations at every observed lymphoid structure, including additional stromal infiltrate regions. Upon data collection the pathologist team annotations were merged with the MIBIscope data by aligning imaging field-of-view (FOV) locations to the H&E image coordinates. If multiple annotated lymphoid structures were captured in a single MIBIscope image, an x or y centroid threshold was used to separate the single cell data into image subsets labeled as -a, -b, or -c and subsequently analyzed as single structures.

### Lymphoid structure numerical scoring

Annotations for each lymphoid structure were converted to numerical values as such: SI=1, LA=2, TLSi=3, TLSm=4. For each structure, the mean and standard deviation of pathologist annotation scores was calculated for each H&E level, as well as the mean across H&E levels termed the “mean maturity score” (MMS). The SD-global was calculated as the mean of the two H&E levels SD,, and the SD-Z was calculated as the difference between the H&E level mean scores.

### MIBIscope imaging

Data was collected at Ionpath Inc (Menlo Park, CA) on their MIBIscope instrument M14. A pathologist-annotated region-of-interest (ROI) H&E map was used to determine the distance (µm) between lymphoid structure centroids, and guide the MIBIscope FOV locations through conversion to motor units, and secondary electron detector images were used to confirm observable tissue architecture to the serial H&E image. For each ROI, a 2x2 tiled set of FOVs were collected. FOV parameters were set at 400x400µm, 1024x1024 pixels, and using the “Fine” setting on the instrument, resulting in a resolution of 0.39um/pixel.

### MIBI image preprocessing

MIBI images were downloaded from the Ionpath MIBItracker cloud platform (https://mibi-share.ionpath.com/) as multi-layer tiffs. MIBItiffs were then preprocessed using the MIBIO software (Ionpath Inc., Menlo Park, CA) to remove bare slide gold adducts, metal ion organic adduct interference (oxide, hydride, and hydroxides), and technical noise according to vendor recommendations. Preprocessed MIBItiffs were then stitched using a rigid stitching script with a 1 pixel empty border between FOVs, and converted to ome.tiff.

### Single cell segmentation and annotation

Cell segmentation was performed on pre-processed, stitched region images using the Cellpose2.0^40^ cyto2 model to segment whole cell boundaries, using a summed signal of VIM, CD45, CD20, CD3, PanCK, CD68, CD14, CD11c, and CD31 as cytoplasmic channels, in addition to the nuclear channel. To improve accuracy, we ingested the Aleynick et al.^41^ human generated cell segmentation dataset, and provided additional in house whole cell boundary annotations in tonsil images, retrained a final whole cell segmentation model using the Cellpose2.0 training GUI. Mean marker intensity (MMI) for all markers was extracted from each whole cell object. Cells were filtered for a minimum nuclear channel MMI threshold, and maximum 181Ta channel threshold, representing nucleated cells within tissue areas, respectively. The MMI of all channels in resultant real cells were then scaled by the mean cell area in the dataset, arcsin transformed, winsorized at 99.9 per marker, and 0-1 normalized. Cell annotation strategy followed seminal MIBI studies^42–44^, using a hierarchical annotation workflow of cellular lineages, then cell types, then cell phenotypes using the FlowSOM algorithm^45^. As detailed in Figure S2, all cells are first clustered into 100 clusters using arcsin-transformed MMI values of all markers excluding the nuclear channel, and major cell lineages are annotated, as well as an “other” class. “Other” cells are then clustered into 25 clusters and clusters with identifiable expression patterns are annotated as the appropriate lineage. Cells of the different major lineages are then isolated and subclustered and annotated into cell types with pathologist review of established marker profiles, select cell types are then clustered into phenotypic subsets.

### Histological Region Masking

Histological region masks corresponding to bare slide, lymphocytic aggregate (“agg”), germinal center, FDC process network, HEV objects, and the tumor microenvironment (“TME”) were generated to define histologic regions of each FOV including the objects using the Artificial neural network (ANN) pixel classifier tool in the QuPath software^46^ (v.5.02) using all channels with raw signal, gradient of magnitude, and hessian max eigenvalue features at the scales of 1, 2, and 4. TME, aggregate, germinal center and bare slide masks were combined into an exported label-tiff image, which was subsequently used in conjunction with single cell centroid coordinates for each region image to determine the cells present in each mask. FDC processes and HEV masks were exported separately, and the total area of each of these combined object masks were quantified per region image.

### Single cell spatial analysis

A pairwise “spatial enrichment” metric was calculated for each cell type pair across a structure region by determining the number of cells of each cell type within a 50μm radius of the seed cell type, averaged over all seed cell type cells across the entire image. In the TME and Agg zones, select cell types and phenotypes were chosen to compute this spatial enrichment metric, based on a priori knowledge of their representation in the two zones. Due to the spatial confines of the Agg zone, a radius of 25μm was used for pairwise spatial enrichment calculation.

TME and Agg Niches were computed from zone-specific pairwise spatial enrichment matrices (calculated above) through the k-means clustering of the matrix into K=9 clusters in the Agg, and K=7 clusters in the TME, using custom python scripts (spatial_enrichment_analysis.py). Niche abundance statistics were evaluated between histological classes using GraphPad Prism (10.6.0).

### Linear discriminant analysis

For training the LDA model, only the SI and GC+ tonsil regions were considered (n=56), representing the extremes of the least and most mature structures. An initial set of 758 features were defined over all regions. In the training set, features were filtered to include only those features that (1) were significantly different between the extreme SI and GC+ tonsil classes (p<=0.05 in a t-test comparison of the two training classes); and (2) had no correlation with any other feature of r>=0.99 on the same training classes. 489/758 features passed these filters. A linear discriminant model^47^ (Ripley) was trained and evaluated for the ability to distinguish SI from GC+ tonsil regions in 5-fold cross validation in the training set at a range of feature numbers. Features were added to the model one-at-a-time in order of the magnitude of the ANOVA F statistic for the two SI and GC+ tonsil classes. The final LDA model was trained with n=489 features, and saved. That trained LDA model was then used to project the full dataset of n=199 regions onto the single linear discriminant axis (LD1), and the distribution along that axis of the other classes, that were not included in training, was examined. Analyses were executed in Python 3.12 using numpy 2.3.2, pandas 2.3.1, and scikit-learn 1.7.1.

### Slingshot analysis

The Slingshot^31^ R package was downloaded from the Github page (https://github.com/kstreet13/slingshot) and deployed on a feature matrix including 759 features defined on 199 ROIs. Features were filtered to those that had a significant difference between the SI and GC+ Tonsil classes (t-test p<=0.01). Features that were very highly correlated with other features (Pearson r>=0.90) were also removed. 355 features remained after those filters. The final feature matrix was standardized so that each feature had zero mean and unit variance. The standardized feature matrix was reduced to two dimensions using UMAP. Using the UMAP projection, regions were clustered into k=5 groups by k-means clustering. Slingshot trajectory analysis was then executed using the UMAP projections, guided by the k-means clusters. Analyses were executed in R 4.5 with Bioconductor 3.21.

### TLS Spatial Network Analysis

To characterize the spatial topology and connectivity of lymphoid environments, we developed a graph theory-based analysis using custom scripts in Python. Centroid coordinates of classified lymphoid structures (LA, LAma, TLSi, and TLSm) were extracted and processed. These centroids served as nodes within spatial graphs constructed using the NetworkX Python library. Undirected edges were generated between nodes based on a proximity threshold; specifically, an edge was established between any two structures located within a 2 mm Euclidean radius of one another—calculated using SciPy (scipy.spatial.distance) representing a biologically relevant interaction distance. Discrete spatial networks were identified as connected components and subsequently stratified based on maturation status: components containing at least one mature TLS were designated as TLSm+ networks, while those lacking mature TLS were classified as TLSm- networks. Network topology was quantified by calculating the node size and edge density for each identified component via NetworkX. To further evaluate the spatial affinity between specific lymphoid aggregate subsets and tertiary lymphoid structures, we performed a nearest-neighbor distance analysis. For each identified LAma and LArest structure, we calculated the minimum Euclidean distance to the nearest TLS (inclusive of both TLSi and TLSm) as well as the mean distance to all TLSs present within the tissue section using SciPy. Statistical differences in network size and connectivity between TLSm+ and TLSm- groups were assessed using the Mann-Whitney test with GraphPad Prism (10.6.0).

### MLS classifier

To identify MLS we designed a classification pipeline using vision-transformer-based embeddings in conjunction with an in-house Detectron2 TLS instance-segmentation model. Briefly, 882x882px crops of the H&E image above each detected LAma and TLS on the MIBi slide were used from annotations for TLSm, TLSi, LAma, LAim, and SI at base-resolution (0.25mpp). These FOVs were then tiled into 98x98px sub-patches, resulting in 81 inputs per FOV being encoded by our internal DINOv2-based ViT L/16^48^. The resulting feature embeddings had hierarchical K-means clustering applied per FOV to determine local morphological groupings. The resulting local clusters from each FOV were then re-clustered to determine global clusters^49^. In order to consolidate each FOV back to a single feature embedding, the cosine distance of each sub-patch embedding is determined relative to the global clusters. The final feature representation for the FOV was a vector of association proportions of each sub-patch to the global clusters (final k=128). A decision tree classifier was trained on the 128-dimensional cluster-based embeddings to classify FOVs as being enriched (TLSm, TLSi, LAma) or unenriched (LA^im^, SI). Application of this classifier to unseen patient whole-slide H&E images first leveraged the TLSv4.1^17^ classifier to identify regions of the H&E image with predicted TLS. Regions that met a probability >0.5 to be a TLS object were captured in a bounding box with the structure at the centroid, and a 882x882px H&E crop was submitted into the workflow as described above. If one or more regions were classified as MLS by the random forest classifier, that patient was identified as “MLS enriched” and accordingly included in the biomarker eligible population for survival analysis.

### Survival Analysis

Survival analysis was performed using the lifelines package50 in Python 3.10 to generate the Kaplan–Meier (KM) survival curves51, and the survival times across the different arms within the IMpower110 clinical trial cohort, and compared using the log-rank test. On the KM survival curve, the vertical and horizontal axes represented the probability of patient survival at a given time. Patients were right censored during the follow-up period if they were event-free, or experienced another non-treatment related outcome. A p-value<0.05 in a two-sided analysis was considered as statistically significant. Hazard ratios (HR) were also calculated to estimate instantaneous risk within Atezolizumab treatment arm when compared to the standard of care arm and evaluate its effect for progression free survival (PFS). If the HR was between 0 and 1, it indicated that the presence of MLS enriched TLS instances was predictive of Atezolizumab efficacy within a given cohort.

## Data Availability

All image data, segmentation data, region masks, single cell feature data, and structure-level summary feature data are available upon request and will be deposited in a public data repository upon acceptance. All code used in the analysis are available upon request and will be deposited in a public code repository upon acceptance.

## Acknowledgements

Specific Contributions: Conceptualization, L.M.M., T.R., R.J.J.; data collection, T.R.; pathology review, L.M.M., N.B., O.L., E.F., L.T., K.P., J.G.; image analysis, T.R., R.J., A.H., C.F.; MLS classifier development, E.L., N.B.; writing T.R. The authors thank Will Ewart and the Genentech Research Pathology Human Tissue lab for the procurement, metadata capture and processing of the study samples, Jian Jiang and the Genentech Research Pathology Histopathology team for the preparation of tissues, and Barzin Nabet, Jeff Eastman, and Robin Lorenz for their editorial input of the manuscript.

## Ethical considerations

The human tissues used to validate this protocol were de-identified, archival specimens acquired from Avaden Biosciences, Bioreclamation, and Cureline Inc.. These specimens were procured in full compliance with HIPAA, IRB protocols, and The Common Rule, with written informed consent for research use obtained by the vendor at the time of collection. Because this protocol utilizes the secondary use of de-identified archival specimens, the requirement for ethical approval by a specific Institutional Review Board (IRB) was waived under US Federal Regulation 45 CFR 46.104(d)(4).

---

**Figure S1:**
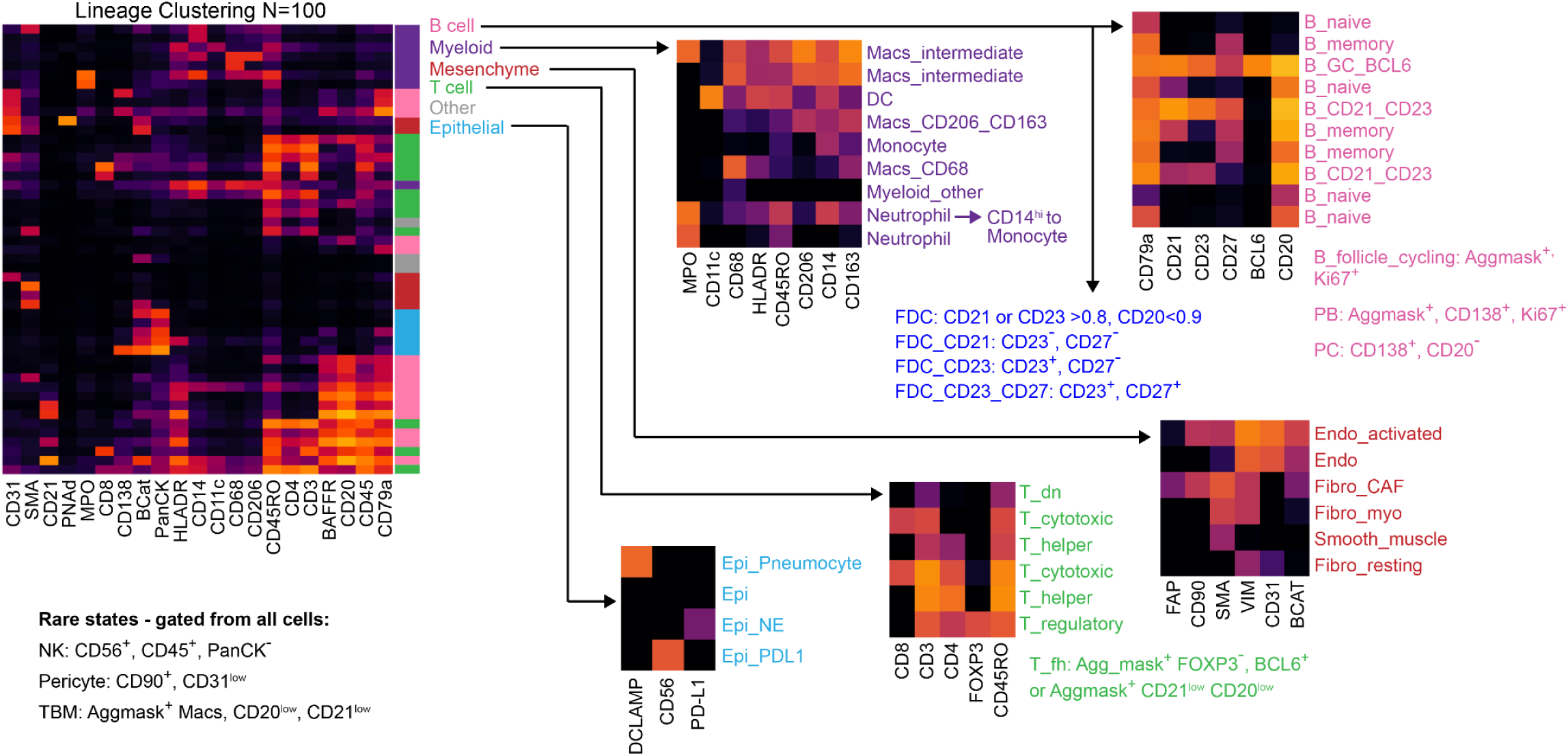
Cell and structure annotation strategies. Schematic showing the hierarchical clustering steps used to annotate single cells in the dataset. The FlowSOM algorithm is leveraged at various steps with unique marker sets to annotate each single cell lineage (left), cell type (middle), and phenotype (right).

**Figure S2:**
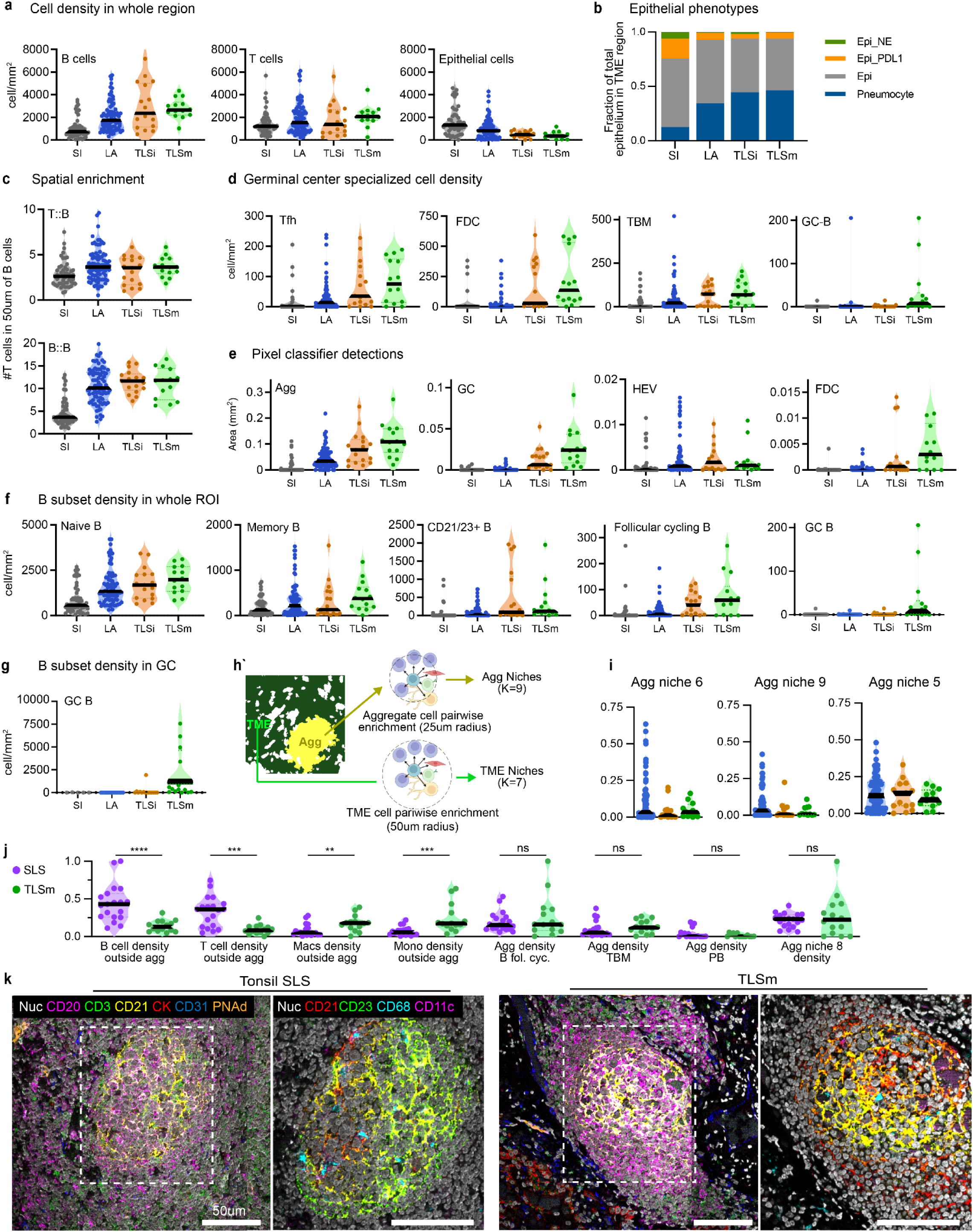
Feature comparison across pathologist classes and tonsil SLS. **a**. Violin plots comparing the single cell density of B cells, T cells, and epithelial cells between histological classes. **b.** Area column plot showing the phenotypic fraction of epithelial cells within each histological class. **c**. Violin plots showing the spatial enrichment of T cells with B cells, and B cells with B cells across the histological classes. **d**. Violin plots showing the density of specialized germinal center related cell types across the four histological classes. **e**. Violin plots showing the area detections of aggregates, germinal centers, HEVs, and the FDC network across the four histological classes. **f**. Violin plot showing the density of B cell phenotypic subsets between the four histological classes. **g**. Violin plot showing the density of GC B cells within the detected germinal center area of each structure, across the four histological classes. **h**. Schematic showing the separation of aggregate and TME cells by region masking and the subsequent niche calculation in the different compartments using distinct radius and cell identify parameters. **i.** Violin plot showing the fraction of aggregate niches between aggregate-positive structures of the LA, TLSi, and TLSm classes. **j**. Violin plot showing the normalized feature level of select features between tonsil SLS and TLSm structures. **k**. MIBI overlays of a representative tonsil SLS and TLSm with major cell type markers to the left, zoomed inset focusing on FDC phenotypic markers CD21 (red), CD23 (green), TBM marker CD68 (cyan), and TBM marker CD11c (magenta).

**Figure S3:**
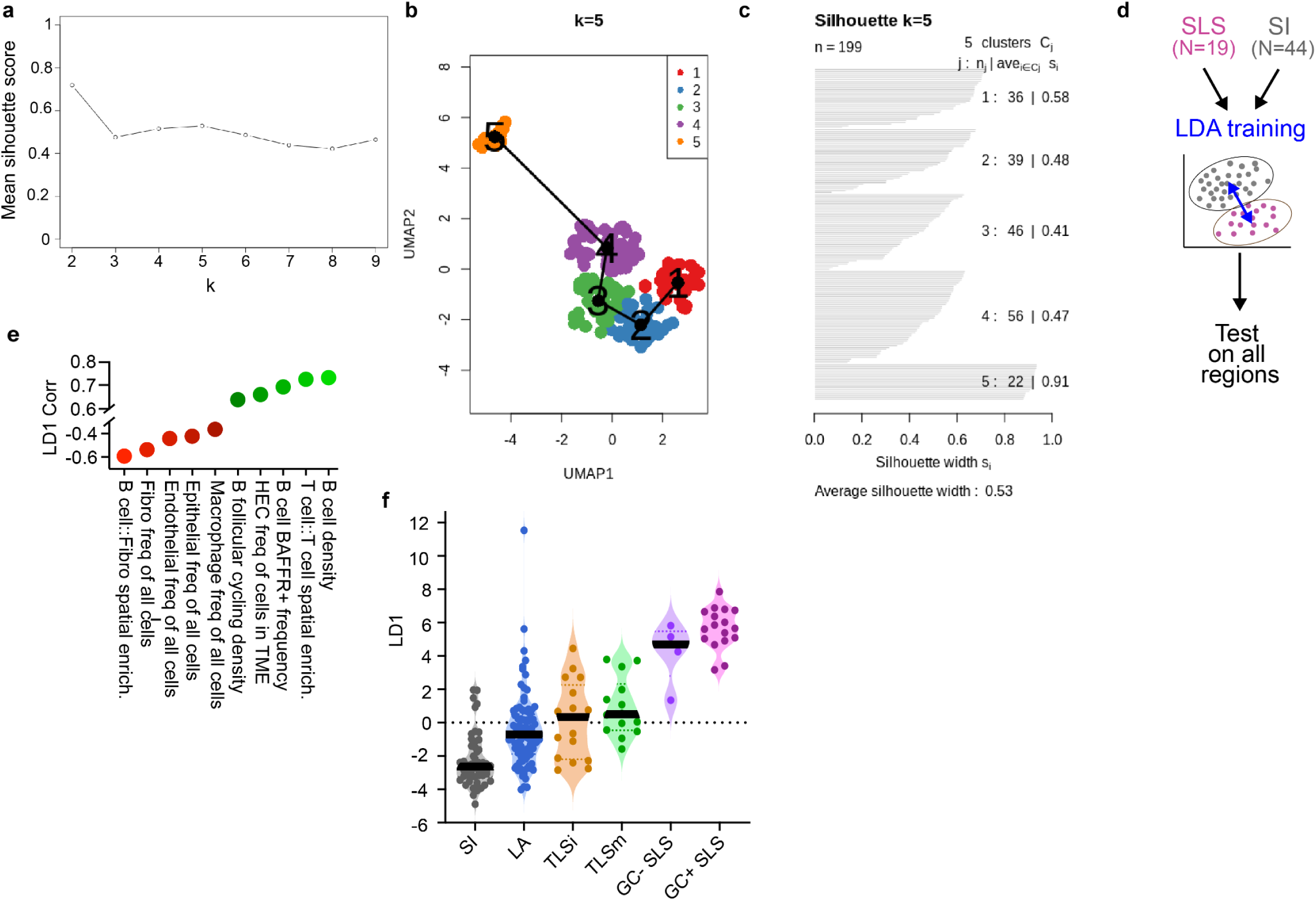
Clustering and classification of tumor lymphoid structures. **a-c**. Silhouette analysis outputs showing the result of different K values on silhouette score using Slingshot feature outputs, with UMAP plot overlays of K cluster identity and silhouette width analysis for each structure. **d**. Schematic showing the structures used to train a LDA classifier to classify unseen structures as either SLS or SI. **e**. Top positive (green) and negative (red) correlated features to the LD1 variable from the LDA classifier analysis. **f**. Violin plot showing the LD1 value for every structure in the cohort, separated by histological class.

**Figure S4:**
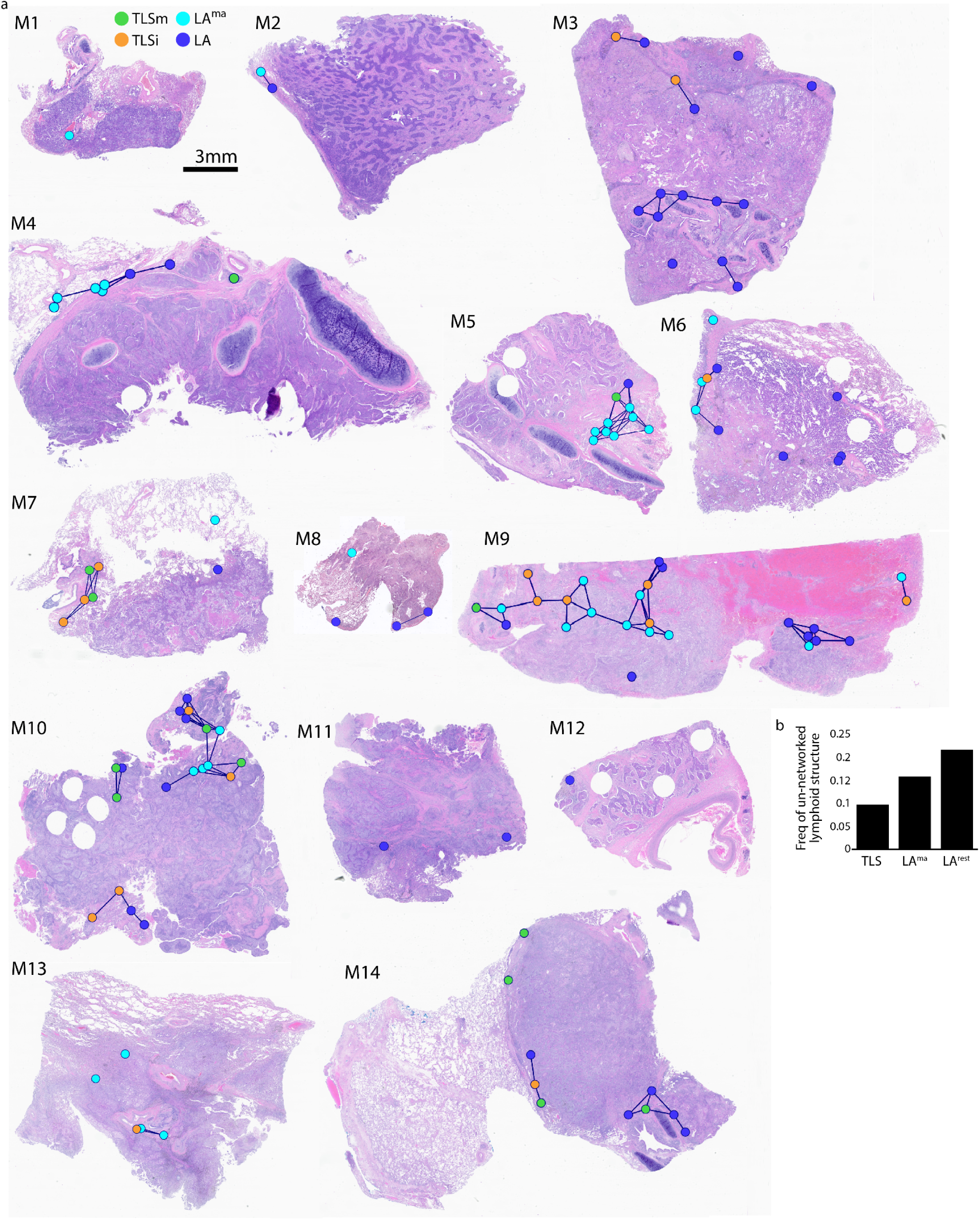
Overview of structure networks across cohort samples. **a**. H&E images of patient tumor resections overlaid with the location of each lymphoid structure, colored by class, including TLSm (green), TLSi (orange), LAma (cyan) and LArest (blue). SI regions not displayed. Structures within networks are connected with black lines denoting network edges.

**Figure S5:**
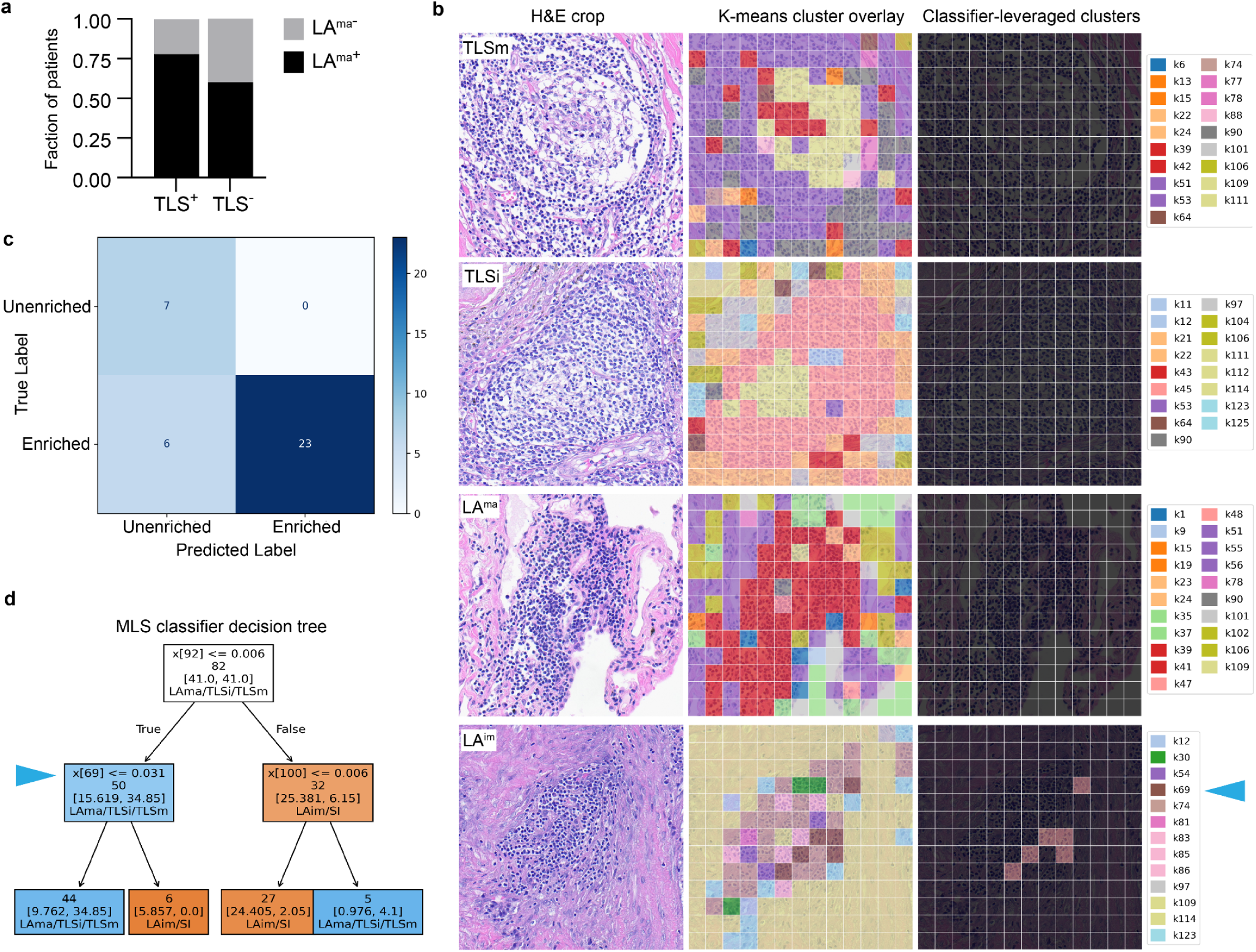
MLS classifier performance and outputs. **a**. Area column plot showing the fraction of samples that contain LA^ma^ between TLS^+^ and TLS^-^ samples. **b.** Representative examples of HE cropped images (left), patch cluster identify (middle) and classifier-leveraged patches (right) from the MLS classifier workflow, for each histological class. **c**. Confusion matrix from in-cohort validation of MLS classifier. d. Random forest decision tree from MLS classifier, with cluster identity denoted as “x[cluster x]”. Blue arrow marks decision point leveraging cluster 69, highlighted in *b*.

